# Gut Microbial Signals Influence Motivational Responses in the Male Mouse

**DOI:** 10.64898/2026.08.21.745917

**Authors:** Brendan L. Sharvin, Gabriel S.S. Tofani, Jade A. Lawless, Lily Keane, Gerard Clarke, Colin G. McNamara, John F. Cryan

## Abstract

**Highlights:**

- Gut microbiome regulates motivation for palatable rewards in a dynamic manner
- Microbial regulation of reward gene pathways in the nucleus accumbens
- Immune, metabolomic and vagal mechanisms interact to drive microbiota-induced stimulation of reward pathways

Mammalian motivation to obtain natural rewards is regulated by internal biological and external environmental factors. Increasing evidence suggests the gut microbiota may represent one such factor. Here, we observed that antibiotic (ABX)-induced disruption of the gut microbiota increased the motivation to obtain palatable rewards, elevated inflammatory cytokine levels and elicited significant alterations in synaptic plasticity-related gene pathways in the mouse nucleus accumbens (NAc). We also observed that the dynamic state of the gut microbiota partially impacts motivational responses and identified several gut microbial metabolites that could be driving this phenotype. Finally, surgical ablation of vagal signalling via subdiaphragmatic vagotomy partially rescued transcriptomic signatures in the NAc and elevations in inflammatory markers but was not enough to prevent the increase in motivational responses. Overall, our results highlight an important new role for the gut microbiota in regulating the cascade of metabolic, neurochemical and immunological events necessary for reward processing and motivation.

Graphical Abstract

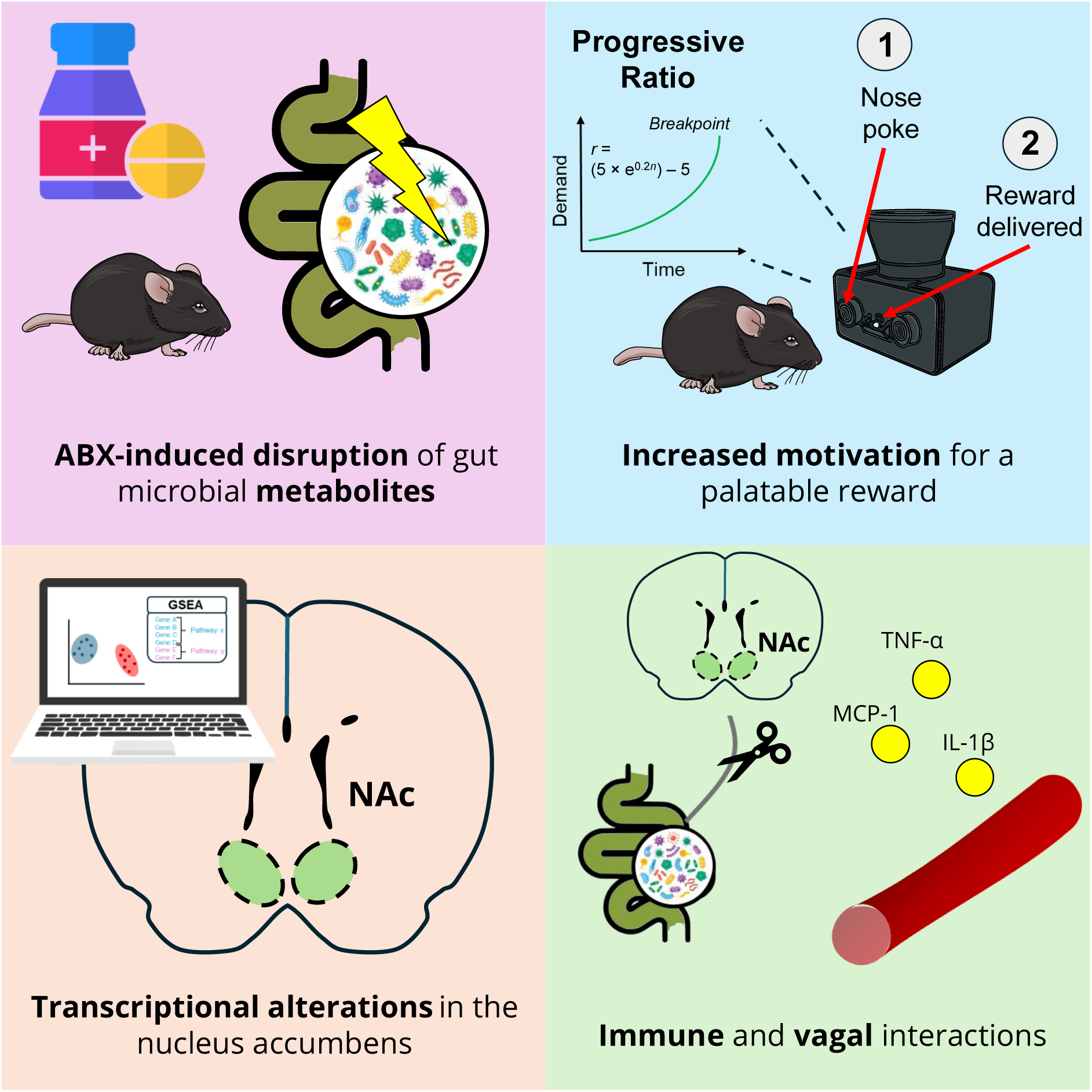

## Introduction

Motivation and decision-making rely on a complex integration of intrinsic and extrinsic factors. A critical component of mammalian survival depends upon the motivation to acquire hedonic rewards such as food, sexual, and social stimuli while interpreting/balancing factors such as demand and choice. These calculations are processed by the mammalian reward system, a complex network of neuronal structures within the brain which allow for the ‘wanting’, ‘liking’ and learning elements of the reward-behavioural cycle (Berridge & Kringelbach, 2015).

Importantly, mammals and microorganisms have co-existed for millions of years throughout adaptations and diversification in a cross species symbiotic relationship (Theis et al., 2016) Indeed, all mammals harbour a gut microbiota and increasing evidence has pointed to a role for these microorganisms in shaping dietary choice and decision-making across numerous mammalian species (Hammer et al., 2019; Mallott et al., 2024; X. Wu et al., 2022). The last decade of research has unveiled how these microorganisms can act as potential regulators of complex higher order behavioural paradigms, including social interaction, fear recall, mood and anxiety levels through a bidirectional communication system referred to as the microbiota-gut-brain axis (Chu et al., 2019; Crumeyrolle-Arias et al., 2014; Cryan et al., 2019; Desbonnet et al., 2013; Hoban et al., 2018; Margolis et al., 2021; Sharvin et al., 2023; W. L. Wu et al., 2021). Furthermore, disrupting gut microbial signals has demonstrated consequences to host feeding behaviour and response to hedonic food reward across multiple species (Mason, 2017; Ousey et al., 2023; Schneider et al., 2024; Trevelline & Kohl, 2022; H. Yang et al., 2018). In parallel, many peripheral cues, which are susceptible to influence from the gut microbiota, such as the immune system and endocrine signals have been well characterised to have a role in modulating reward processes (Ben-Shaanan et al., 2016; Boyle et al., 2023; Cassidy & Tong, 2017; Greene et al., 2019; Montesinos et al., 2016). However, the neurochemical mechanisms that could be responsible for driving these behavioural phenotypes remains poorly understood.

There is evidence to suggest that such behavioural adaptations induced by gut microbial signals could be mediated via vagal afferent fibres which innervate enteroendocrine cells along the gastrointestinal (GI) tract and terminate in the nucleus tractus solitarius (NTS; Fülling et al., 2019). From here, an inter-synaptic pathway, involving multiple brain nuclei, interact to induce dopamine release in the striatum (Han et al., 2018).

At the forefront of reward neurocircuitry is the nucleus accumbens (NAc), a subcortical brain structure located deep within the frontal lobe and a hub for interpreting cues about hedonic rewards, pleasure, and motivation through processing of dopaminergic projections from the ventral tegmental area (VTA). Indeed, there is some existing evidence to suggest a role for microbial signals in influencing this critical brain region (García-Cabrerizo et al., 2021, 2024). Germ free (GF) rodents were shown to have significant alterations in neuronal morphology within the NAc, specifically dendritic elongation of medium spiny neurons (MSNs) in the core and dendritic hypertrophy in the shell (García-Cabrerizo et al., 2024). Moreover, ABX-treated female mice displayed altered cocaine reward processing, which corresponded with distinct dopamine and immediate early gene-related transcriptional changes in the NAc (Dave et al., 2025). This suggests gut microbial signals may have a role in programming normal development and neuronal activity of the NAc.

Taken together, there is evidence of the gut microbiota’s role in regulating motivation for palatable reward, influencing neuronal morphology in the NAc, however a clear mechanism connecting these findings remains unknown. Thus, our goal in this study was to understand the relationship between the gut microbiota and the motivation to obtain palatable rewards and assess the potential signalling mechanisms, including the vagus nerve underwriting such effects.

## Results

### Microbial depletion with antibiotics elicited an increased motivation to obtain a palatable reward

To address the role of the gut microbiota in directing motivational responses to hedonic palatable rewards we treated mice with antibiotics (ABX) to disrupt the gut microbiota and employed an operant conditioning paradigm (Figure 1A). During this protocol, mice were food restricted, and importantly we observed no significant differences in weight throughout this process of ABX-treated animals compared to controls (mixed effects model (treatment): F (1, 18) = 2.415; p = 0.1376; Figure S1A). Mice were first trained to learn the nose poke-reward association. As control and ABX-treated mice then advanced from a fixed ratio (FR) of one nose poke per reward (FR1), to three (FR3) and then six pokes (FR6) their discrimination between the active and inactive nose port developed (mixed effects model (nose poke); control: F (1, 18) = 69.34; p < 0.0001; ABX: F (1, 18) = 111.7; p < 0.0001). Moreover, both groups gradually increased the number of active nose poke entries across sessions ((nose poke x session) control: F (8, 138) = 12.48; p < 0.0001; ABX F (8, 142) = 15.49; p < 0.0001; Figure 1B and C). Strikingly, when comparing groups, ABX-treated mice outperformed control mice and earned significantly more pellets throughout the training period (mixed effects model (treatment): F (1, 18) = 29.77; p < 0.0001), independent of the increasing phases of demand ((treatment x session): F (8, 139) = 1.426; p = 0.1607). Following this FR training period, mice advanced to a progressive ratio (PR) task, where the amount of nose pokes required to earn a single pellet increased exponentially, at which point mice reach a “breakpoint” (the maximum number of nose pokes or physical demand required to earn a reward before a subject ceases to engage). The breakpoint is considered a measure of the animal’s motivation levels (Cambre et al., 2023; Hailwood et al., 2018; Rivera et al., 2025). Interestingly, ABX-treated mice elicited a significantly higher breakpoint in the PR task, both under food restriction (unpaired t-test; p = 0.011) and when food was available *ad libitum* (unpaired t-test; p = 0.004) when compared to controls (Figure 1E; see Table S1 for additional statistical information). Importantly, when measuring chow intake per cage, there were no significant differences in regular chow consumption between ABX and control cages (n = 3 / group) during the operant task, when food was returned to *ad libitum* availability, suggesting effects on motivation were not due to satiety levels (mixed effects model (treatment): F (1, 4) = 0.4903; p = 0.5224; Figure 1F). Overall, this suggests that gut microbial disruption with ABX resulted in an increase in effort to obtain a palatable reward across a FR training schedule, and during the PR task, suggesting an increase in motivational responses.

**Figure 1.**
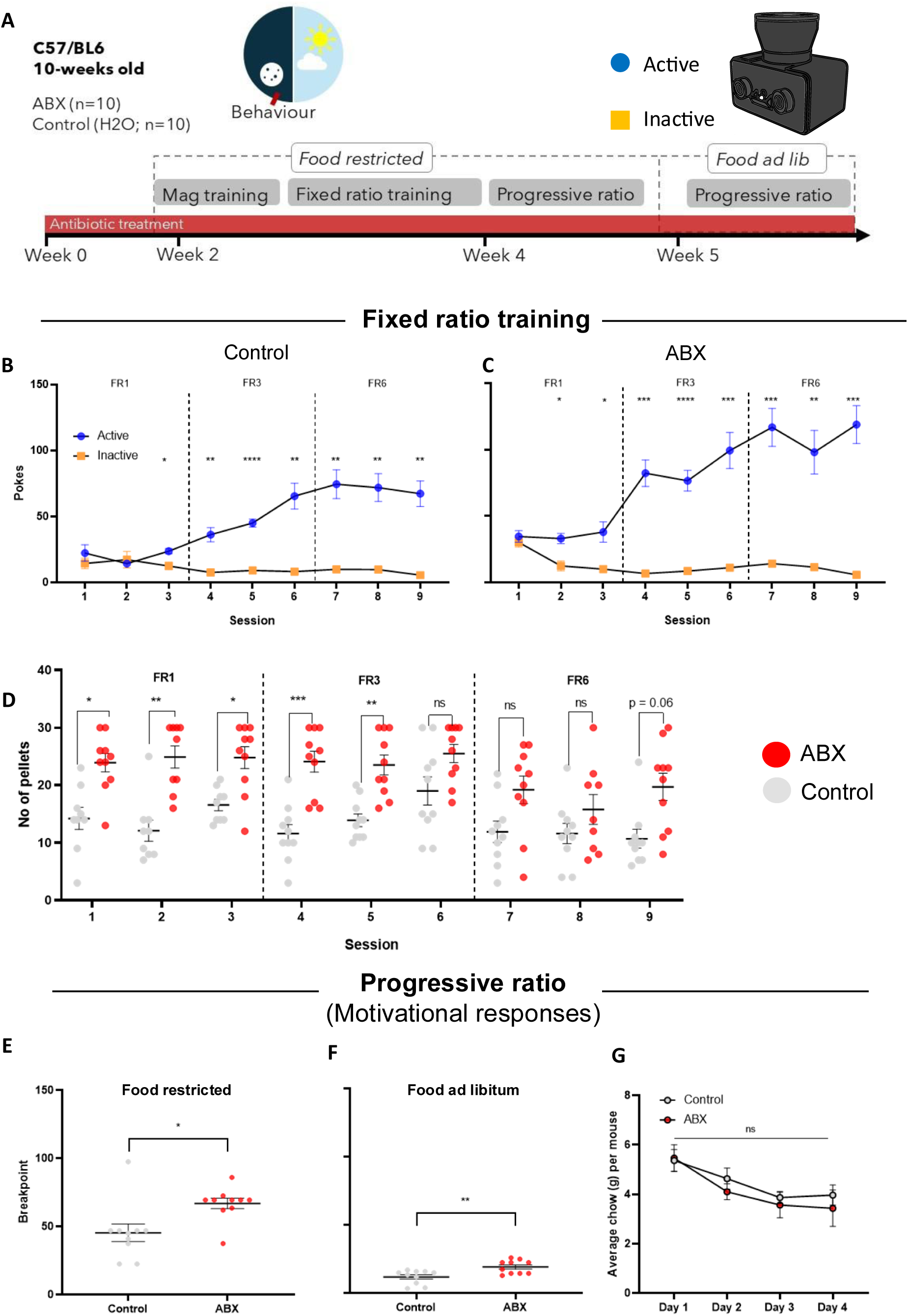
ABX results in a significant increase in effort to obtain a palatable reward in an operant conditioning protocol. (A) Experimental protocol and operant training schedule for control (n = 10) and ABX (n = 10) mice. (B) Number of active and inactive nose pokes (mean +/- SEM) of control mice across the FR1, FR3 and FR6 operant training sessions. (C) Active and inactive nose pokes (mean +/- SEM) of ABX mice across the FR1, FR3 and FR6 behavioural sessions. (D) ABX mice earned significantly more palatable rewards across the FR training protocol (30-minute sessions). (E) ABX mice elicited a higher breakpoint in the PR task both when food restricted and when (F) food was available ad libitum. (G) No significant differences in average chow consumption (g) per cage (n = 3 per group) between ABX and controls, when food was returned to ad libitum. Data presented as mean +/- standard error of mean (SEM). (B-D, G) Mixed effects models with repeated measures followed by Sidak’s multiple comparisons test. (E, F) PR breakpoints assessed via unpaired t test. * p<0.05, ** p<0.01, *** p <0.001, **** p < 0.0001 and ns = not significant. Detailed statistical information can be found in Table S1.

### ABX results in significant alterations to gene pathways in the nucleus accumbens relevant to dendritic spine, ribosomal structure and the extracellular matrix

Following the discovery that ABX treatment resulted in a significant increase in motivation to obtain a palatable reward, we sought to understand the neurochemical consequences of disrupting gut microbial signalling on reward processing neurochemistry. To address this, we performed RNA sequencing of the nucleus accumbens (NAc; Figure 2A). Initially, we performed principal component analysis (PCA) and conducted a PERMANOVA to assess transcriptomic differences between control and ABX on a compositional level (Figure 2B) and observed no significant changes (see Table S1 for additional statistical information). Following this, we employed a gene set enrichment analysis (GSEA; using the gseGO function from clusterProfiler) to identify gene pathways in the NAc that are affected by ABX treatment using the Gene Ontology database (Ashburner et al., 2000; Figure 2C). Interestingly, we observed a significant upregulation of pathways relevant to ribosomal DNA transcription and the dendritic spine (Figure 2C). Dendritic morphology in the NAc has previously been linked to the gut microbiota, as well as dictating motivation levels in rodents (Bayassi-Jakowicka et al., 2021; Bringas et al., 2013; García-Cabrerizo et al., 2024). Indeed, we also observed significant downregulation of pathways relevant to the extracellular matrix in the NAc (Figure 2C). In the same animals, we also assessed systemic pro-inflammatory cytokine levels in the blood (Figure 2D). Interestingly, we observed an increase in TNF-α, MCP-1, and IL-1β (one-way ANOVA with Tukey’s multiple comparisons test (control vs ABX); TNF-α (p = 0.025), MCP-1 (p = 0.031), and IL-1β (p = 0.087); see Table S1 for additional details of statistical information) following ABX treatment (Figure 2E-G). Taken together, these transcriptomic shifts in synaptic plasticity-related pathways in the NAc could be leading to altered reward processing following a disruption to typical microbial-gut-brain communication. Furthermore, the alteration in certain peripheral inflammatory cytokines suggests inflammatory signals may be a potential signalling mechanism driving this phenotype (Ben-Shaanan et al., 2016; Boyle et al., 2023; Huwart et al., 2025).

**Figure 2.**
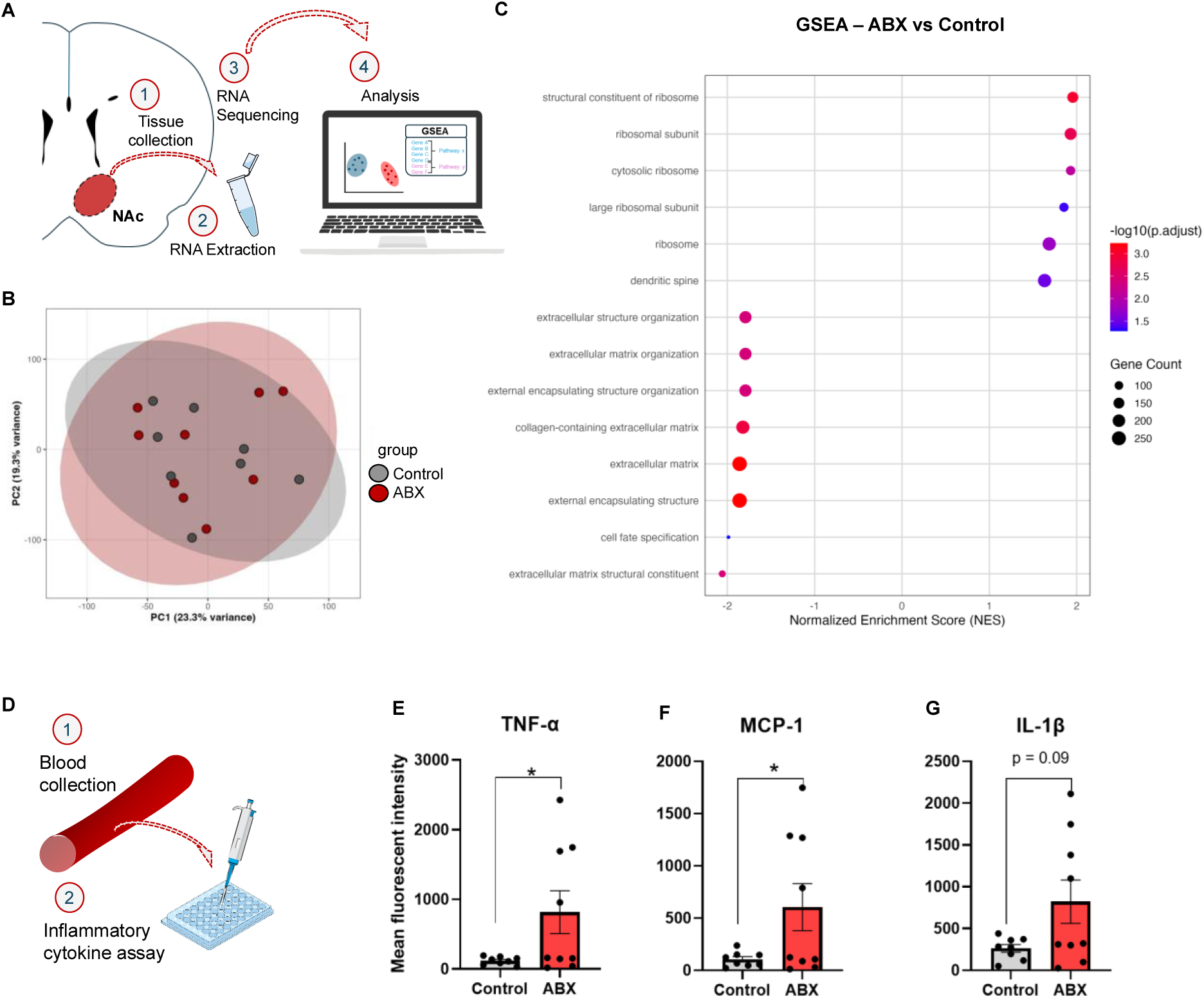
ABX-induced depletion of the gut microbiota leads to significant alterations in gene pathways in the NAc and increased systemic inflammatory cytokines. (A) Schematic workflow of RNA-Seq of the NAc. (B) Principal component analysis of the NAc transcriptome (Control, n=8; ABX, n=9). Data analysed by PERMANOVA (C) Gene set enrichment analysis of NAc transcriptome showing pathways upregulated (NES (+)) and downregulated (NES (-)) following ABX treatment. Enrichment plots showing ABX enriched pathways with p-adjusted < 0.1. (D) Schematic workflow of systemic inflammatory cytokine assay. (E-H) Increase in mean fluorescent intensity of the pro-inflammatory cytokines in the serum for (E) TNF-α, (F) MCP-1 and (G) IL-1β following ABX treatment. Data analysed by one-way ANOVA, followed by Tukey’s multiple comparisons tests. * p<0.05, ** p<0.01, *** p <0.001, **** p < 0.0001 and ns = not significant. Detailed statistical information can be found in Table S1.

### The dynamic state of the gut microbiota partially dictates motivation for palatable reward

Next, we wanted to further comprehend the longitudinal impact on motivational responses (PR breakpoints) following dynamic disruptions to gut microbiota composition and gut-brain signalling. We established two new experimental groups, ABX-washout and ABX-delayed groups which underwent a similar operant conditioning protocol. The ABX-washout group received ABX during FR training before removal of ABX whereas the ABX-delayed group acquired the task and were then placed under ABX treatment. As mice progressed from FR1 through FR3 to FR6, these mice too increased the nose pokes and discrimination for the active port (mixed effects model (nose poke): ABX-delayed; F (1, 10) = 7.883; p = 0.0185; ABX-washout; F (1, 8) = 5.680; p = 0.043). Again, during FR training schedule, mice that received ABX showed trends to earn more pellets (ABX phase: F (1, 9) = 3.996; p = 0.0767; see Figure S2). Following recording of baseline PR breakpoints for both groups, mice performed behavioural sessions in block schedules comprised of FR6 (engagement) and PR sessions (motivational responses; Figure 3 and S2). ABX-delayed mice that achieved a baseline level of motivation (Block 1), elicited a significant increase in motivation for palatable reward across sessions (mixed effects model (sessions): F (2.262, 11.31) = 16.24; p = 0.0004: Figure 3B and C). When comparing PR breakpoints in ABX-delayed mice from block 1 and block 4, there was a significant increase in motivational responses (p = 0.0175). Interestingly, ABX-washout showed no effect of washout across sessions on motivational responses (sessions: F (1.332, 5.329) = 3.251; p = 0.1252; Figure 3D and E)). Furthermore, when comparing block 1 against block 4, PR breakpoints in ABX-washout mice showed no differences (p = 0.9678). This suggests that this behavioural phenotype is partially dictated by the dynamic state of the gut microbiota and recolonisation of the ABX-depleted gut microbiota failed to recover motivational responses.

**Figure 3.**
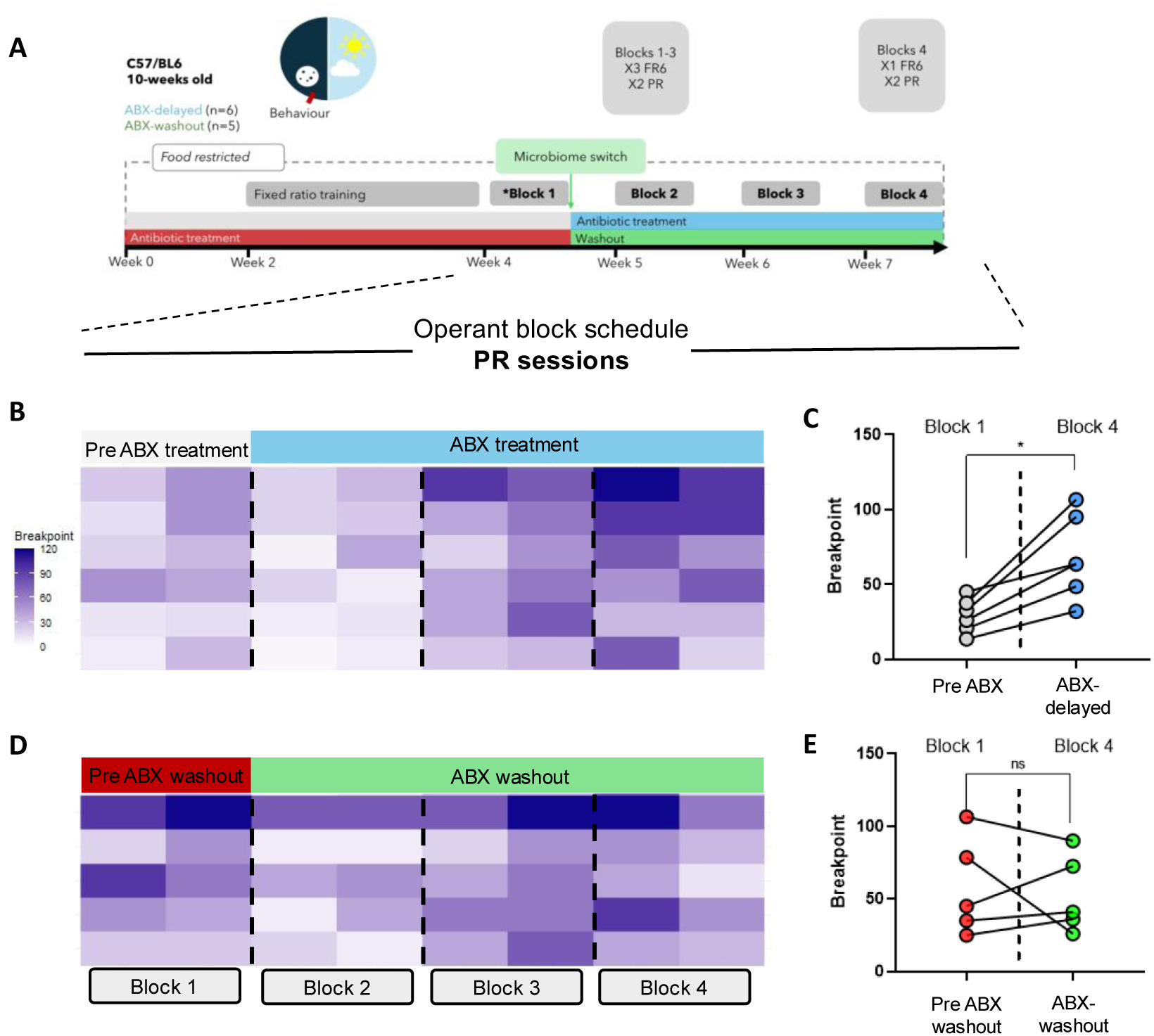
The dynamic state of the gut microbiota partially impacts metrics of motivation for palatable reward. (A) Experimental protocol with operant training schedule, microbiome switch and block schedules. (B) Heatmap of within-subject breakpoints across 4 block operant schedules for pre- and post-ABX-treated mice (ABX-delayed group; n=6). (C) Before-after dot plot of average breakpoints with connecting lines showing within-subject motivation between blocks 1 and 4. (D) Heatmap of within-subject breakpoints across 4 block operant schedules for pre- and post-ABX-washout mice (ABX-washout group; n=5). (E) Before-after dot plot of average breakpoints with connecting lines showing within-subject motivation between blocks 1 and 4. (C/E) Data presented as average (mean) breakpoint across 2 sessions within a block. Data analysed by mixed effects models with repeated measures followed by Dunnett’s multiple comparisons test. ∗p < 0.05, ∗∗p < 0.01, ∗∗∗p < 0.001, ns = not significant. Detailed statistical information can be found in Table S1.

### ABX induced significant restructuring of the caecal metabolome which was partially recovered following washout

Next, we sought to understand how this shift in microbiota composition following ABX treatment could impact the metabolic profile of the caecum. Specifically, we wanted to understand the metabolites and/ or pathways that were altered in ABX mice, stratify based on gut microbial-associated metabolites and identify a potential mechanism through which the gut microbiota could be mediating motivational responses. To achieve this, we performed untargeted metabolomic analysis of the caecal content in control, ABX, ABX-washout and ABX-delayed mice. Furthermore, given the role of short chain fatty acids in brain and behaviour (O’Riordan et al., 2022) we performed a targeted SCFA analysis of caecal contents. Following PCA, we observed differences in degrees of variance between our experimental groups (Figure 4A; see Table S1 for statistical information). This drastic restructuring in the caecal metabolome is illustrated in a volcano plot showing metabolites differentially expressed in ABX compared to control mice (Figure 4B). Of the significantly altered metabolites, we selected the top 40 based on effect size in the ABX mice (overlap with ABX-delayed mice, see Figure S3). Of these 40 metabolites, we conducted a literature search to identify the top gut-microbial associated metabolites (highlighted in green in volcano plots; see Table 1). We conducted a pathway enrichment analysis and found ABX mice were enriched in pathways relevant to phenylalanine, tyrosine and tryptophan biosynthesis, along with many others (Figure 4C). Many of these pathways have functional relevance to feeding neurocircuitry (Castells-Nobau et al., 2024; Green, 2025; Zhu et al., 2024). Given the finding, that ABX-washout mice maintained these atypical increases in motivational responses, we wanted to identify what metabolites failed to recover or persisted to increase following this washout period. Interestingly, we observed that most gut microbial associated caecal metabolites recovered, apart from indole-3-propionic acid and taurochenodeoxycholate-3-sulphate (Figure 4D). This is further illustrated in the heatmap comparing log fold change of these metabolites in each group relative to the control (Figure 4E). Indole-3-propionic acid is a metabolite derived from tryptophan by the gut microbiota. Recent evidence supports a role for this metabolite in affecting neuronal activity, potentially providing neuroprotection and raising kynurenic acid levels in the rodent brain (Owe-Larsson et al., 2025; Sathyasaikumar et al., 2024). There is little evidence to show taurodeoxycholic-3-sulphate has effects on brain function and behaviour via gut-brain signalling, however its role in influencing immune response suggests the immune system could be a potential indirect signalling mechanism influencing this behavioural phenotype (Xu et al., 2025). Unsurprisingly, we also observed a significant reduction of SCFAs in the caecum following ABX treatment (Figure 4F). Interestingly, ABX-washout failed to recover levels of 2-methylpropanoic acid, propanoic acid (or proprionate), acetic acid and pentanoic acid to levels comparable to controls (see Table S2). Interestingly, delivery of propionate has previously been shown to reduce reward response in the human striatum (Byrne et al., 2016). Indeed, we also observed subtle trends to suggest acetic acid, propanoic acid, and pentanoic acid displayed negative correlations with motivation for palatable reward (see Table 2). Taken together, untargeted and targeted metabolomic analyses revealed a broad restructuring of the caecal metabolome following antibiotic treatment, including significant shifts in microbial-associated metabolites and enrichment of pathways linked to aromatic amino acid biosynthesis. Notably, several gut-derived metabolites and key short-chain fatty acids failed to recover after washout—most prominently indole-3-propionic acid, taurochenodeoxycholate-3-sulphate, and multiple SCFAs—highlighting potential microbiota-dependent mechanisms are dysregulated.

**Figure 4.**
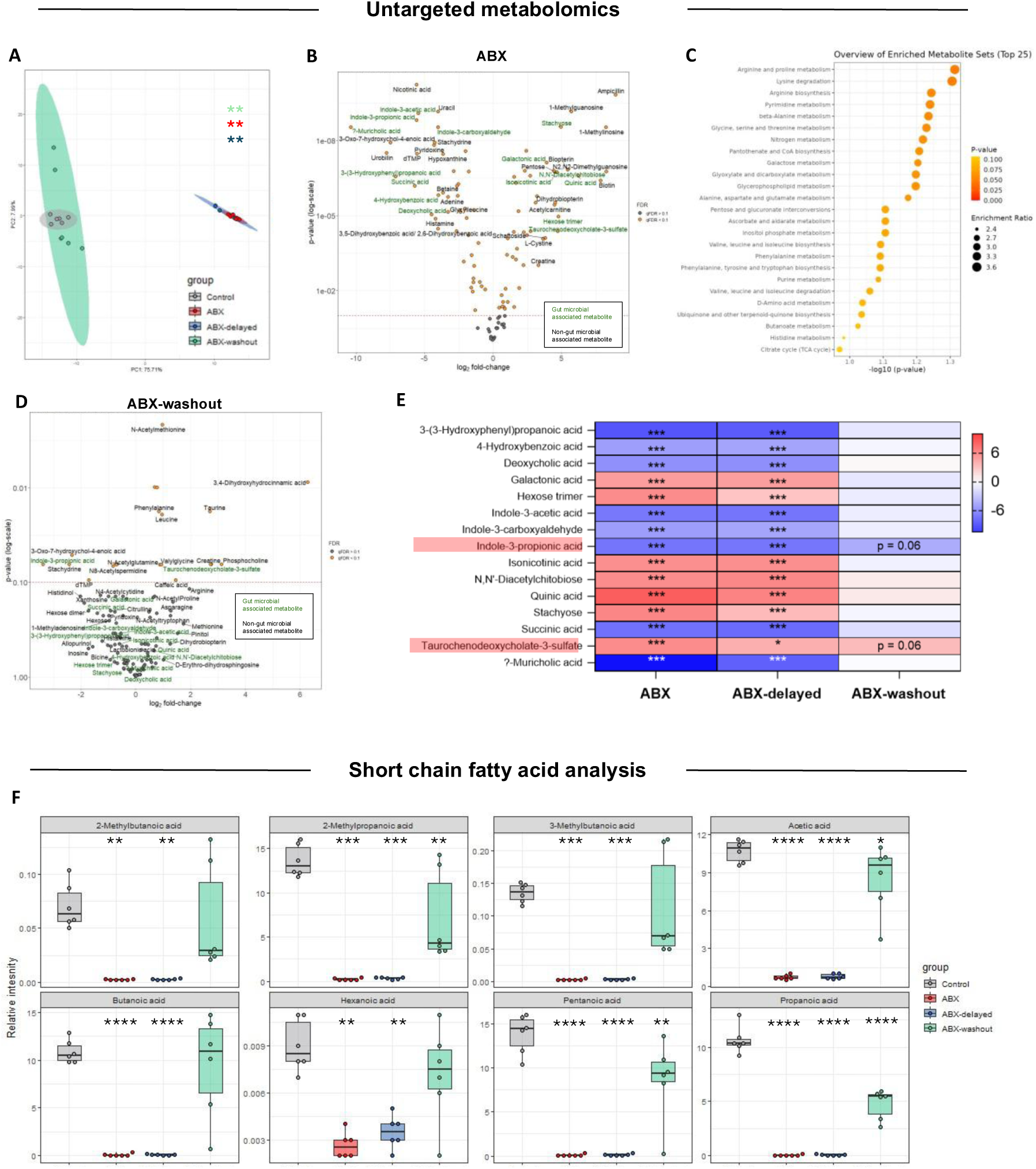
ABX and ABX-washout resulted in a significant restructure of the caecal metabolome compared to conventional controls. (A) PCA plot showing variance across 4 experimental groups (n = 6/ group). Data analysed by PERMANOVA. (B) Volcano plot showing significantly altered metabolites and their log-fold change in ABX compared to control mice, with multiple testing correction (q<0.1; n = 6/ group)). (C) Enrichment analysis showing metabolites enriched in ABX compared to control (following FDR correction, q < 0.1; n = 6/ group)). (D) Volcano plot showing significantly altered metabolites and their log-fold change in ABX-washout compared to control mice, with multiple testing correction (q<0.1; n = 6/ group)). (E) Heatmap showing log2 fold change of top gut microbial-associated metabolites across each group relative to control (n = 6/ group). (F) Boxplots of relative intensity values of SCFAs following targeted SCFA analysis that are significantly different between Control and ABX, ABX-delayed and ABX-washout (n = 6/ group). Data presented as mean +/- SEM. Untargeted metabolomics data analysed using PERMANOVA with 1000 permutations and Benjamini-Hochberg post-hoc test to control for FDR. Targeted SCFA analysis data analysed using one-way ANOVA with Dunnett’s multiple comparisons test. ∗p < 0.05, ∗∗p < 0.01, ∗∗∗p < 0.001, ns = not significant. Detailed statistical information can be found in Table S1.

**Table 1.**
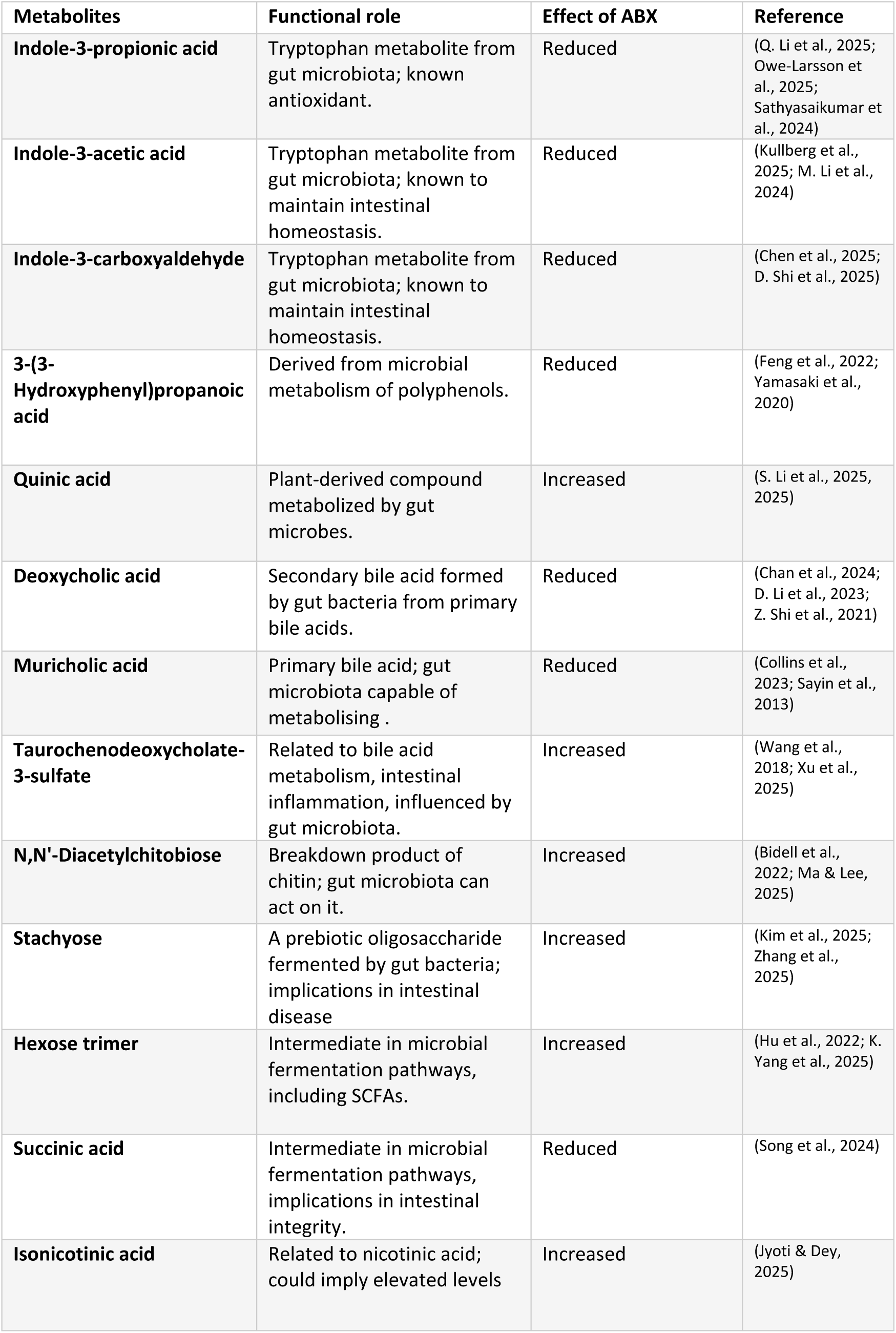
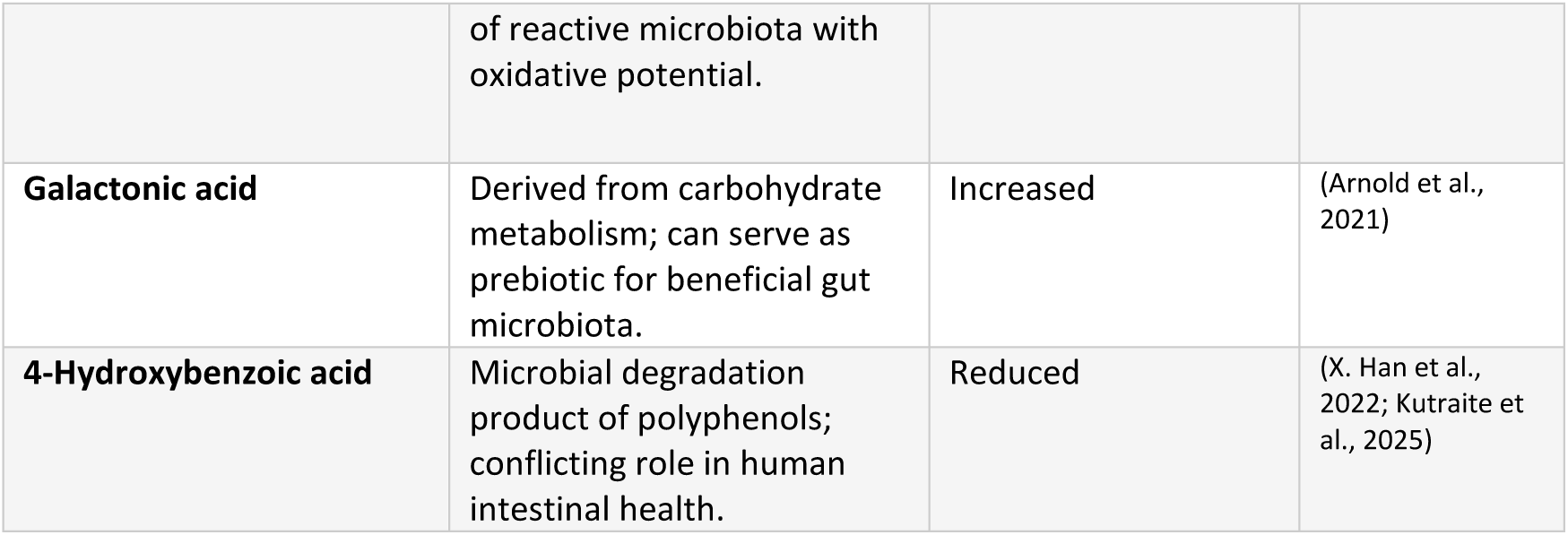
Gut microbial-associated metabolites dysregulated following ABX treatment (from untargeted analysis Fig 4B).

**Table 2.** Correlation of log-transformed peak area of SCFAs (from targeted analysis Fig 4F) against scaled breakpoint in PR task. Data analysed by Spearman’s correlation, with FDR correction (p.adj).

| Metabolite | rho | p.value | p.adj | Correlation |
| --- | --- | --- | --- | --- |
| **2-Methylpentanoic acid** | 0.394447 | 0.062526 | 0.203208 | Positive |
| **Acetic acid** | -0.40682 | 0.05404 | 0.203208 | Negative |
| **Hexanoic acid** | -0.48304 | 0.019555 | 0.203208 | Negative |
| **Pentanoic acid** | -0.40583 | 0.054683 | 0.203208 | Negative |
| **Butanoic acid** | -0.2913 | 0.177478 | 0.329602 | Negative |
| **Heptanoic acid** | -0.30223 | 0.161025 | 0.329602 | Negative |
| **Propanoic acid** | -0.31784 | 0.139417 | 0.329602 | Negative |
| **2-Methylbutanoic acid** | -0.18193 | 0.40607 | 0.52789 | Negative |
| **2-Methylpropanoic acid** | -0.18344 | 0.402135 | 0.52789 | Negative |
| **4-Methylpentanoic acid** | 0.194167 | 0.374674 | 0.52789 | Negative |
| **3-Methylbutanoic acid** | -0.16642 | 0.447884 | 0.529317 | Negative |
| **Formic acid** | -0.13099 | 0.551332 | 0.597276 | Negative |
| **3-Methylpentanoic acid** | -0.09952 | 0.651434 | 0.651434 | Negative |

### Subdiaphragmatic vagotomy partially prevents ABX-driven transcriptional changes in the NAc and peripheral cytokines, yet produces limited improvements in ABX-associated motivational impairments

The vagus nerve forms an integral part of mediating gut-induced alterations to brain and behaviour (Bravo et al., 2011; Breit et al., 2018; Fülling et al., 2019) and has been shown to mediate gut-induced reward processing in the striatum (W. Han et al., 2018; Kaelberer et al., 2018). Thus, we next hypothesised that the vagus nerve was a key signalling mechanism which could be driving this ABX-induced elevation in motivational responses and subsequent disruptions to gene pathways in the NAc. To address this, we performed subdiaphragmatic vagotomy (SDV) to surgically ablate vagal afferents prior to ABX treatment protocol and trained mice as previously described (Figure 5A). Importantly, there were no significant differences in initial ABX drinking water consumption following SDV when compared to controls (mixed effects model (vagus status); F(1, 18) = 1.205; p = 0.2867; Figure S4F). Importantly, all groups increased the number of nose pokes and discrimination for the active port as they progressed throughout the FR learning paradigm (see Table S2 for additional statistical information; Figure S4A-D). Interestingly, the ABX-induced increase in effort to obtain palatable reward throughout the FR learning phase persisted (mixed effects model (treatment): F(1, 33.98) = 27.06; p < 0.0001) independent of vagal signalling (vagus status: F(1, 33.98) = 0.004; p = 0.947; Figure 5B; S4A-D; see Table S1 for additional statistical information). Furthermore, the ABX phenotype on motivation for reward in the PR task prevailed (two-way ANOVA (treatment); F (1, 31) = 5.122; p = 0.0308) independent of vagal signalling (vagus status: F (1, 31) = 1.477; p = 0.2333; Figure 5C). Despite this, we did observe that SDV rescued the ABX-induced alterations observed in the NAc transcriptome, specifically the dendritic spine pathway, alterations to ribosomal function and downregulation of the extracellular matrix network (Figure 5D). To further explore this, we filtered total genes based on gene pathways enriched in the SHAM-ABX (Figure 2C) and plotted the top 50 genes based on effect size and observed that SDV indeed rescued many of the ABX-induced alterations in individual genes driving dendritic spine, ribosomal function and extracellular structure (Figure 5E). This implies that perhaps these ABX-induced alterations to reward processing we previously identified in the NAc are vagal-dependant, at least partially. Given the link between the gut microbiota and the immune system, we measured inflammatory responses in the SDV model. Interestingly, the increased inflammatory response we observed in ABX treated mice appeared to be absent in the SDV-ABX animals (Figure 5F-H; see Table S1 for additional statistical information), indicating that the vagus nerve is also necessary in mediating ABX-induced disruption to immune signalling. To assess whether these alterations in palatable reward were driven by alterations in glucose sensitivity, we measured glucose levels in the blood and found no effect of ABX (two-way ANOVA (treatment); F(F (1, 34) = 0.3389; p = 0.5643) nor SDV (vagus status: F (1, 34) = 0.1477; p = 0.4281; Figure S4G). Taken together, SDV reversed ABX-induced transcriptional and immune disruptions in the nucleus accumbens, however the heightened motivational drive for palatable reward appeared to persist independent of vagal signalling. This suggests a complex integration of multiple signalling mechanisms could be driving this ABX-induced phenotype on motivational responses.

**Figure 5.**
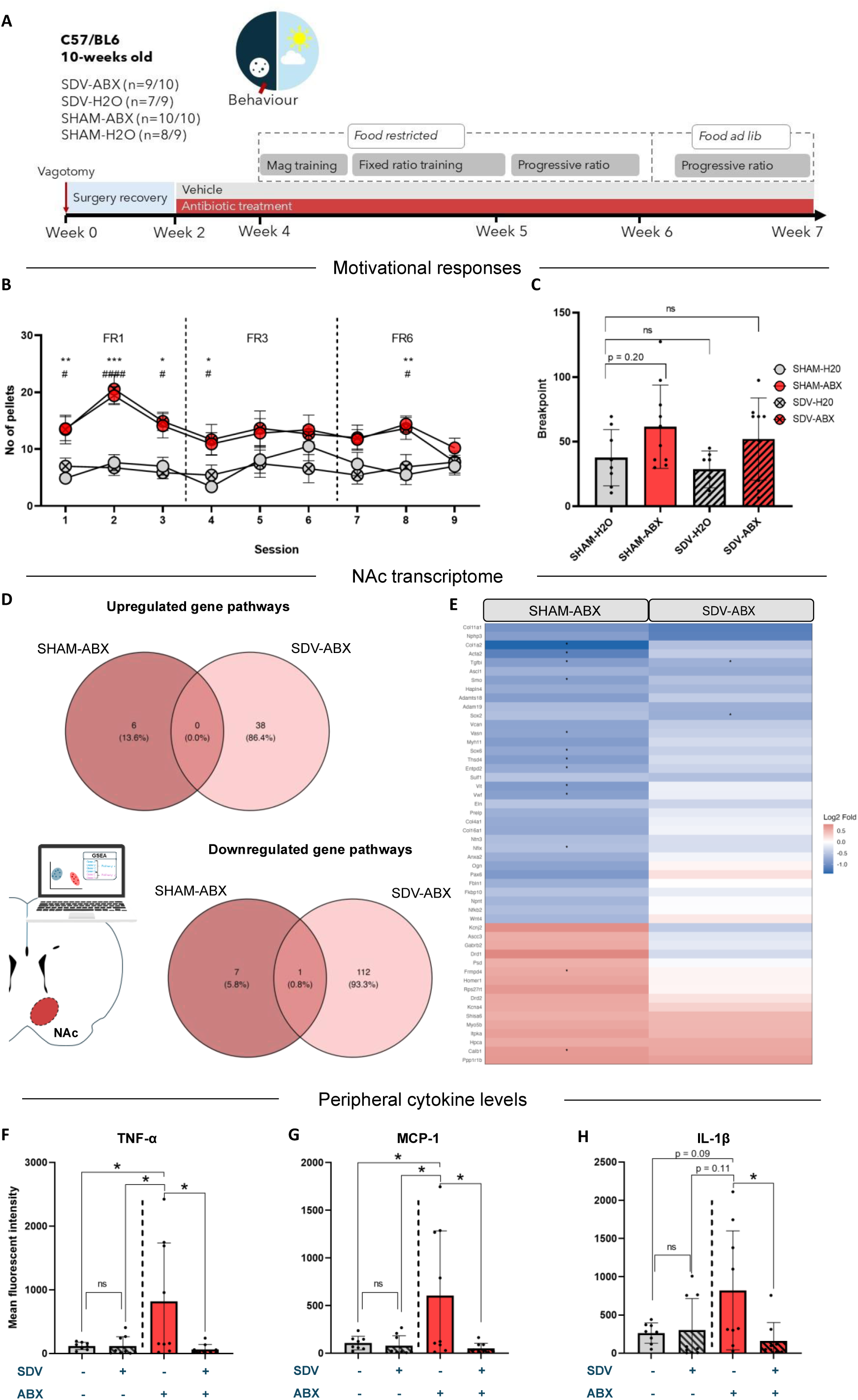
Surgical ablation of vagal signalling prevents ABX-induced alterations to NAc gene pathways, increases in inflammatory cytokines, but not motivational responses. (A) Experimental protocol showing subdiaphragmatic vagotomy (SDV), subsequent recovery period and operant training schedule. (B) The number of pellets earned across FR1, FR3 and FR6 training schedules (30-minute sessions). Data analysed by mixed models (treatment, vagus status and session as factors), with post-hoc multiple comparisons * SHAM-H2O vs SHAM-ABX, # SHAM-H2O vs SDV-ABX. (C) Progressive ratio average breakpoint across 2 sessions showing effect of ABX, but no effect of SDV. Data analysed by two-way ANOVA (treatment, vagus status as factors). (D) Venn diagrams showing upregulated and downregulated pathways induced by ABX treatment appear to be vagal dependant (overlap of only 1 pathway: cell fate specification). (E) Heatmap of genes related to pathways altered by SHAM-ABX, filtered by effects size (top 50) showing log fold change against of SHAM-ABX and SDV-ABX against controls (SHAM-H20). Raw p values represented by stars. (F-H) Inflammatory cytokine levels (TNF-α, MCP-1, and IL-1β) in the serum from systemic blood showing effect of SDV and ABX treatment. Control data taken from Figure 2 E-G. Data analysed by one-way ANOVA with Tukey’s multiple comparisons test. Data presented as mean +/- standard error of mean. SHAM-ABX n=10, SHAM-H2O n=8/9, SDV-H2O n=7/9, SDV-ABX n=9/10 successfully trained mice. ∗p < 0.05, ∗∗p < 0.01, ∗∗∗p < 0.001, ns = not significant. Detailed statistical information can be found in Table S1.

## Discussion

Understanding the communication pathways between the gut microbiota and reward processes is key to unravelling key survival mechanisms in mammals. Taken together, this research has identified a role for the gut microbiota and subsequent microbial signalling in mediating intrinsic motivation for hedonic palatable rewards, NAc transcriptome changes and peripheral immune signalling coupled with a restructuring of the caecal metabolome and altered production of microbial metabolites.

Our findings further expands upon previous work investigating this intrinsic role of gut microbial signals in mediating response to palatable (Ousey et al., 2023) and other rewards (Berland et al., 2022; Dave et al., 2025; Dohnalová et al., 2022; García-Cabrerizo et al., 2021; Kiraly et al., 2016). Importantly, here we revealed that ABX-induced depletion of the gut microbiota increased the motivation to obtain palatable rewards along with alterations to synaptic plasticity related gene pathways in the NAc. Moreover, partial faecal microbiota transfer and ABX-washout failed to recover metrics of motivation to levels comparable to controls. Given this finding, we wanted to identify whether we captured a persistent perturbation of the gut microbial profile that could be attributed to this atypical increase in motivation. Following analysis of the caecal metabolome, we identified several gut microbial associated metabolites that failed to recover/normalise, including indole-3-propanoic acid, propionate, and acetic acid.

Interestingly, oral propionate supplementation has previously been shown through fMRI recordings to reduce anticipatory reward responses to high energy foods in the human striatum (Byrne et al., 2016). Thus, a reduction in propionate levels following ABX, and subsequent failure to recover in the ABX-washout mice could explain this persistent elevated motivation for palatable rewards and transcriptomic alterations in the NAc. Indeed, ABX-induced depletion of the gut microbiota has been shown to enhance cocaine seeking behaviour in an operant task, along with altered transcriptional alterations in the NAc, which were reversed with SCFA supplementation, further emphasizing the role of the gut microbiota and metabolites in mediating mesocorticolimbic networks and reward seeking behaviour (Acuña & Olive, 2024).

At the level of the NAc, we observed significant changes in gene pathways that could be attributed to motivational responses following ABX treatment, including dendritic spine, ribosomal transcription and the extracellular matrix. In general, ribosomal activity in the NAc is modulated by external stimuli, such as exposure to drugs. In fact, previous research has shown that increases in ribosomal DNA transcription in MSNs within the NAc were identified in heroin addict patients (Gos et al., 2022). An increase in dendritic spine density and morphology was observed in rodents following chronic social defeat stress (Fox et al., 2020). Indeed, such stressors have also been shown to mediate effort-based reward driven motivation in mice models (Evertse et al., 2024). Moreover, dendritic hypertrophy and morphological alterations were observed in the NAc of GF mice (García-Cabrerizo et al., 2024). Furthermore, ABX has been shown to elicit altered dendritic morphology in other brain regions including the hippocampus and somatosensory cortex (Nakhal et al., 2025). Alterations in neuronal morphology within the NAc, including the dendritic spine, have previously been shown to elicit alterations in reward processing and motivation levels (Colom et al., 2024; Fox et al., 2020; Hikida et al., 2016). The extracellular matrix provides a scaffold, maintaining the architecture of neuronal networks. Disruptions to the extracellular matrix in the NAc are characteristic of disrupted synaptic plasticity, altered neuronal excitability and weakened perineuronal nets, thereby potentially affecting motivational responses to reward stimuli (Hikida et al., 2016; Mews et al., 2024). Taken together, these transcriptomic alterations in gene pathways following ABX-induced disruption to microbial-gut-brain signalling may be contributing to enhanced reward response via disrupted synaptic plasticity and extracellular architecture. Importantly, we have not explored the role of microbial signals and metabolites in mediating dopamine dynamics in brain reward areas in response to palatable rewards, and future investigations should consider applying *in vivo* recording techniques such as fiber photometry or optogenetics to untangle these mechanisms.

We also observed significant elevation in systemic inflammatory markers in ABX-treated mice. Depletion of the gut microbiota has previously been shown to induce effects on the immune system, shifting systemic inflammatory markers in the blood (Fülling et al., 2020; Lynch et al., 2023; O’Riordan et al., 2025; Ratsika et al., 2023; Takiishi et al., 2017). The observed increase in peripheral inflammatory cytokines raises the possibility that immune activation contributes to the motivational phenotype observed in ABX-treated animals. Although we cannot assign causality, the combined molecular and behavioural alterations point toward a broader immune-to-brain communication axis that may shape motivational processes under conditions of microbiota disruption. Interesingly, alterations in the adaptive immune system have been shown to affect motivation for palatable reward and subsequent reward processing in the brain (Boyle et al., 2023; Felger & Miller, 2012; Huwart et al., 2025).

Despite the well-established role of the vagus nerve in directing microbial-gut-brain signaling, the neurochemical consequences of ABX-induced depletion of the gut microbiota with and without the absence of vagal input remains relatively elusive. Here, we demonstrated for the first time, to the best of our knowledge, that depletion of the gut microbiota via ABX triggers vagal-dependent immune and transcriptomic changes in reward circuitry whilst sparing motivational behavioural responses per se. Specifically, we observed that the ABX-induced increase in inflammatory cytokines was vagal dependant. The vagus nerve has previously been identified to mediate systemic immune responses via the cholinergic anti-inflammatory pathway (Alen, 2022; Liu et al., 2025). It is plausible that ABX-induced disruption to the gut microbiota alters this signalling and leads to a systemic pro-inflammatory response which requires interactions with vagal afferents to trigger a central neuroimmune response in the brain. Moreover, we observed that the ABX-induced alterations to the dendritic spine, extracellular matrix and ribosomal pathways were dependant on the vagus nerve. This was further illustrated in the heatmap where we subsetted our RNA-Seq dataset based on these pathways and plotted the top 50 genes based on effect size. Despite the ABX-induced alterations to the NAc transcriptome being mediated via vagal signals, we observed that the effect on motivational responses persisted. This important discovery suggests alternative signalling routes may be reinforcing this ABX-driven behavioural phenotype. For instance, the involvement of spinal afferents, independent of vagal signalling, has been shown to mediate gut microbial reward response to exercise in rodents (Dohnalová et al., 2022). We also cannot rule out the possibility that the systemic shifts in inflammatory markers and circulating metabolites may enter the brain via the blood brain barrier. Previous research has linked the gut microbiota to changes in barrier integrity both in the brain and gut (Braniste et al., 2014; Hoyles et al., 2021; D. Shi et al., 2025; Takiishi et al., 2017).

### Limitations of the study

This research was conducted on males only. The justification was two-fold: studies have shown stronger microbiota mediated effects on brain and behaviour in males, and there are clear sex-specific effects on motivational responses to palatable reward (Grimm et al., 2025; Kumar et al., 2024; Reichelt et al., 2016; Clarke et al., 2013; Geary et al., 2021; Jaggar et al., 2020; Jašarević et al., 2016). Nevertheless, for future studies it is important to decipher sex-dependant differences following these findings.

It is important to note that the sample size power calculation for our follow-up experiment which included the ABX-delayed and ABX-washout group yielded relatively low numbers and was based on the large effect size on behaviour we observed in our initial study comparing ABX and controls. It is also possible that a larger sample size would further uncover a more nuanced role for the vagus nerve in mediating this ABX-induced phenotype. Furthermore, it is possible that the exposure to condensed milk (liquid diet) during the recovery phase following surgery may have influenced our results.

We did not investigate the effects of psych biotics that might counter the effects of ABX-induced microbiota depletion or modify these behaviours when the gut microbiota is fully intact. Indeed, it has been shown previously that ABX-treated mice consumed more palatable rewards and this phenotype was rescued via colonisation of bacterial taxa from the S24-7 family and *Lactobacillus johnsonii* (Ousey et al., 2023).

## Conclusion

Taken together, our data further supports the role of the microbiota in driving mammalian motivational responses. The divergent effects of vagotomy on microbiota depletion-induced-changes in inflammatory response and reward processing further expands our understanding of microbial-immune crosstalk and brain-body interactions. The persistence of increased motivational responses for palatable rewards in the absence of typical vagal communication highlights the complexity of microbiota-gut-brain axis signalling and the involvement of multiple signalling mechanisms, including microbial metabolites and immune processes. Overall, these findings suggest a potential interoceptive dysregulation of typical motivational processes following a removal of gut microbial signals, which may have broader implications across neuropsychiatric processes from anhedonia to addiction phenotypes.

## Resource availability

### Lead Contact

Further information and requests for resources and reagents should be directed to and will be fulfilled by the lead contact, Professor John F. Cryan at.

### Materials availability

This study did not generate new unique reagents.

## Supporting information

Training criteria and engagement

## Acknowledgments

The author would like to thank Patrick Fitzgerald, Ken O’ Riordan, Gerard Moloney, Colette Manley, Naomi Gavioli, Bella Ziade, Linda Katona, Melanie Depret, Ruben Garcia Cabrerizo, and Benjamin Valderrama for their technical assistance and support throughout the project. The research was conducted in the APC Microbiome Ireland, which is funded by Science Foundation Ireland (<u>SFI/12/RC/2273_P2</u>). This research was funded by the Saks Kavanaugh Foundation.

## Author contributions

Conceptualization, B.L.S., C.M., G.S.S.T. and J.F.C.; methodology, B.L.S., C.M., G.S.S.T. and J.F.C.; investigation, B.L.S., C.M., G.S.S.T., L.K., and J.F.C.; data curation, B.L.S., G.S.S.T., L.K., and J.L.; validation, C.M, G.S.S.T and J.F.C.; writing – original draft, B.L.S.; writing – review & editing, B.L.S., C.M., G.S.S.T., J.L., G.C., L.K., and J.F.C; funding acquisition, J.F.C; resources, C.M., J.F.C.; and supervision, C.M. and J.F.C.. All authors contributed to the article and approved the final manuscript.

## Experimental Model and Subject Details

### Studies in Animals

All procedures were conducted with approval from the Animal Experimentation Ethics Committee (AEEC) at University College Cork and the Health Products Regulatory Authority (HPRA) under the project authorizations: AE19130/P188, in accordance with the recommendations of the European Directive 2010/63/EU. C57BL/6 male mice (8-weeks) were ordered from Envigo (https://www.inotiv.com/horizon-welcome). The mice were kept in a holding room with 12:12-h reversed light/dark cycle (lights off at 9am and on at 9pm), controlled temperature and humidity (20+/- 1 degree Celsius, 55.5%). Mice were moved to cages of 2-4 and had 1 week to acclimatize to the holding room before procedures began. Animals were provided with food and water *ad libitum* initially. Prior to operant training, mice were food restricted to 2-3g of chow (maintained at >85% of baseline body weight) which was provided after each daily behavioural session. As the behavioural experiment involved sucrose (palatable) reward, this ensured mice would not be fully satiated and therefore more motivated to engage in the operant tasks.

## Method details

### Antibiotic treatment

Mice were treated with an antibiotic cocktail (ABX) comprised of Ampicillin (1ug/mL), Vancomycin (0.5ug/mL), Gentamicin (1ug/mL) and Imipinem (0.25ug/mL). These antibiotics were administered in drinking water. Fluid intake was monitored, the cocktail was prepared every 2 days and weight was recorded daily for the first 2 weeks of treatment.

### Subdiaphragmatic vagotomy

All surgeries were performed under aseptic conditions following the guidelines outlined by the Laboratory Animal Science Association (LASA; https://lasa.co.uk/wp-content/uploads/2025/03/Guiding-principles-on-good-practice-for-Animal-Welfare-and-Ethical-Review-Bodies-2015-PDF-1.76MB.pdf). Mice were acclimatised to liquid diet (condensed milk mixed with water) 2-3 days prior to the subdiaphragmatic vagotomy surgery. 24 hours before surgery, mice were food deprived. On the day of surgery, mice were anesthetized with isoflurane (5% induction, 1.5-1.8% maintenance). The pedal reflex was assessed before surgery to ensure an appropriate level of anaesthesia was reached. Mice were administered a subcutaneous dose of meloxicam (5mg/kg) and eye gel was applied. Mice were then placed on a sterile gauze on a warm heating blanket. Fur was shaved from the abdomen. The abdomen was then cleaned with sterile 70% ethanol and iodine solution. Sterile autoclaved instruments were used to cut a 2-4cm incision roughly 0.5 cm below the sternum. The skin and peritoneum were separated using forceps. Following midline laparotomy, the stomach was located and externalized. The subdiaphragmatic vagus nerve was revealed by gentle retraction of the stomach and oesophagus. Sutures were fed underneath the gastroesophageal junction to expose the oesophagus. The right and left branches were dissected using curved forceps under the microscope. Sham surgery mice underwent the same protocol, except the subdiaphragmatic vague nerve was spared. Organs were kept moist with regular cleansing with sterile saline. The peritoneum was sutured using vicryl absorbable sutures, while the external skin was sutured using prolene sutures. A total of 47 mice underwent surgery (SDV: 27, Sham: 20). Of the 27 mice that underwent SDV, 3 died under anaesthesia, and 5 developed gastroparesis (delayed stomach emptying) and were euthanised. Of the 20 Sham mice, 1 recovered poorly from surgery and was euthanised.

Mice were left to recover single housed and received meloxicam (5mg/kg) mixed with the liquid diet (condensed milk) for 2-3 days. Following this, mice were weaned to a solid food diet for a further 3 days and received solid diet for the duration of the experiment. The success of SDV was verified by ex vivo observation.

### Food restriction

To maintain motivation during the operant task for palatable reward, mice were food restricted to 3g of chow/ day. Body weight was recorded every day and maintained at a level of >85% of baseline (weight at day before food restriction). Animals were provided with food daily at the early stages of the dark phase, following any behavioural task that took place that day. Ad libitum water access was provided throughout the duration of the experiments.

### Partial faecal microbiota transfer

To allow an ABX-washout in our experiments, on the day ABX-treated mice ceased treatment, faecal material and bedding material from control counterparts were mixed in the cage of the ABX-washout mice to promote recolonisation of the microbiota (Bidot et al., 2018; Budden et al., 2024; Clavell-Sansalvador et al., 2024; Gregor et al., 2020).

### Operant conditioning

Operant conditioning was carried out using the open-source battery-powered Feeding Experimentation Device version 3 (FED3.1; Matikainen-Ankney et al., 2021). The FED3 devices were placed in a cage with a 3-D printed cover. Before each behavioural session, the devices were turned on, and the desired program was selected using the nose pokes. Once the session was selected, the device and cover were placed into cages. These cages were identical to home cages, except without enrichment materials. Clear cage tops were used to facilitate recording from a camera placed above the cages. Sessions lasted between 15 minutes and 1 hour. After each session, mice were returned to their home cage, and the devices and covers were cleaned with 70% ethanol. The screen on the side of the device displays the number of active and inactive nose pokes and the number of pellets distributed. These were manually recorded immediately after the session by the researcher.

#### Magazine training

Mice were habituated to the operant cage and FED3 device for 15 min and then returned to their home cage once a day, for at least 4 days. During these sessions, pellets were freely available in the central magazine of the FED3 device. Removal of a pellet from the magazine would automatically trigger the release of another.

#### Fixed ratio training

Following magazine training, mice were trained on a fixed ratio (FR) training schedule, whereby a nose poke into the active port of the FED3 device would trigger the release of a pellet (FR1). Any nose pokes into the inactive port would be recorded but not trigger the pellet release. On the first day of FR1 training, the active nose port was primed with a palatable pellet to encourage mice to poke in the active port over the inactive port. Following active nose poke entry and delivery of reward, further nosepoke responding without a previous entry into the magazine would fail to trigger the delivery of another reward. Mice were trained on this FR1 schedule (sessions between 15-30 mins). Successfully trained mice were defined as having obtained at least a single reward for 3 consecutive days during FR1 training schedule. Successfully trained mice advanced to the increasing demand sessions. Mice who did not successfully meet the training criteria were removed from the operant protocol, but tissue was collected for processing at end of experiment. Following FR1 training, mice advanced to an FR3 schedule where the demand for a reward increased to 3 nose pokes into the active port. Finally, mice advanced to an FR6 schedule, whereby 6 nose pokes into the active port were required for delivery of a pellet. The location of the active nose port remained consistent for each mouse but counterbalanced across experimental subjects. Successfully trained mice advanced to the progressive ratio (PR) task.

#### Progressive ratio

Following FR training, mice advanced to a PR paradigm. The objective is to assess the animal’s ability to engage in effort to obtain a reward while facing a progressively increasing demand. To receive a single pellet reward, the amount of nose pokes required followed an exponential schedule, *r* = (5 × e^0.2*n*^) – 5, rounded to the nearest integer, with *n* as the position in the sequence of ratios. The last level completed before time out was considered the breakpoint of the animal. PR sessions lasted for 60 mins. PR sessions were conducted every second day, with FR sessions conducted every other day, to maintain the animal’s engagement with the behavioural task. PR sessions were conducted both under food restriction and when food was available ad libitum. Data for PR sessions are presented as an average of 2 repeated session breakpoints.

### Operant block schedules

Following a full operant schedule, any mice that continued operant procedures underwent behavioural assessment in block schedules. The goal of these blocks was to assess longitudinal metrics of motivation (i.e, PR breakpoint). These block sessions were conducted weekly and comprised of daily sessions of either FR6 or PR. PR sessions took place either side of an FR6 session. Again, data for PR sessions are presented as an average of 2 repeated session breakpoints. As PR breakpoints were metrics of motivation, subjects were given a minimum engagement threshold and those that failed to earn at least 1 reward across all FR6 sessions in that block (low-effort benchmark) were removed from longitudinal assessment.

### Fresh tissue collection

Mice were transferred to a separate cull room and euthanised via decapitation. Trunk blood was collected into K2 EDTA lavender-top vacutainers (BD Life Sciences). Following collection, the blood was centrifuged at 4000g for 15 mins at 4°C. Tissues were harvested and stored in PCR-grade 1.5ml tubes and were immediately flash-frozen in dry ice. All tissue samples were stored at −80°C. To collect fresh frozen nucleus accumbens (NAc) tissue, a brain matrix was used to punch out fresh tissue, using the Allen Brain Atlas as a reference (https://mouse.brain-map.org/static/atlas).

### RNA extractions

RNA from the NAc was extracted using the RNeasy Extraction Kit (Qiagen) according to the manufacturer’s instructions. To lysis the tissue, zirconia/ silica beads (Biospec) were used. RNA quality and concentration was then calculated using Nanodrop Spectophtometer (Nanodrop technologies).

### RNA-sequencing

mRNA sequencing (RNA-Seq) was conducted by Azenta (ultra-low input RNA-Seq). Reference genome of obtained sequences was performed using the following reference annotation: Mus musculus (organism), GRCm38, UCSC, genome browser (GRCm38.p6), Ensemble.

### RNA-sequencing data processing, differential expression and gene set enrichment analysis

Raw paired-end FASTQ files from mouse brain RNA-seq experiments were processed using the nf-core/rnaseq pipeline (v3.19.0; Ewels et al., 2020), which performs adapter trimming, quality control, and transcript quantification. All parameters were run with default settings (Ewels et al., 2020), except that pseudo-alignment was enabled via Kallisto (--pseudo_aligner kallisto) and full alignment was skipped (--skip_alignment true). Reads were quantified against the Mus musculus GRCm39 reference genome with Ensembl gene annotations (release 113).

Transcript-level abundance estimates (HDF5 format) were imported into R (v4.5.1) via RStudio (v2025.05.1+513) using the tximport package (v1.36.1; Soneson et al., 2015) for gene-level summarisation. Genes with ≤10 counts in ≥90% of samples were excluded from analysis. Counts were centre log-ratio (CLR) transformed, and principal component analysis (PCA) was performed using the decostand function from the vegan package (v2.7.1; Oksanen et al., 2012). Four outlier samples were identified and excluded based on PCA. The SHAM-H₂O group was used as the reference level throughout. Group-level differences in gene expression were assessed via permutational multivariate analysis of variance (PERMANOVA) using the adonis2 function (1,000 permutations; formula: ∼ group) from the vegan package (Oksanen et al., 2012).

Differential gene expression analysis was performed using DESeq2 (v1.48.2; Love et al., 2014) with the model ∼group and default parameters. P-values were adjusted using the Benjamini– Hochberg procedure to control the false discovery rate (FDR; Benjamini & Hochberg, 1995) genes with adjusted p < 0.05 were considered differentially expressed.

Gene ontology (GO) annotations for detected Ensembl gene IDs were retrieved using biomaRt (v2.64.0) with the mmusculus_gene_ensembl dataset. Genes were ranked by log₂ fold change (log₂FC), and gene set enrichment analysis (GSEA) was performed using the gseGO function from clusterProfiler (v4.16.0). GO terms with adjusted p < 0.1 were considered significantly enriched.

To assess pathway-level biological relevance, we identified *a priori* in the Gene Ontology database (Ashburner et al., 2000; Supplementary Table; file name: “GO_functions_of_interest.csv”) to filter GSEA results. Enriched pathways were classified as upregulated or downregulated based on the sign of the enrichment score, and overlapping pathways between ABX groups were visualised using ggvenn (v0.1.19). Enrichment plots displaying normalised enrichment scores (NES) were generated for significantly enriched GO terms in the SHAM comparisons.

To examine functional convergence, genes annotated to the significantly enriched GO terms in the SHAM-ABX group were analysed across all groups relative to SHAM-H₂O and visualised in a heatmap. Genes with raw p < 0.05 were considered differentially expressed. All visualisations were generated in R using ggplot2 (Ginestet, 2011), unless stated otherwise.

### Inflammatory cytokine assay

Inflammatory cytokine levels in serum were determined using the Legendplex mouse inflammation 13-plex panel, (Biolegend, Cat. No. 740446); a bead-based assay that allows simultaneous measurement of analytes based on cell size using flow cytometry. Specifically, the cytokines measured were CCL2 (MCP-1), GM-CSF, IFN-β, IFN-γ, IL-1α, IL-1β, IL-6, IL-10, IL-12 (p70), IL-17A, IL-23, IL-27 and TNF-α. The assay was carried out in a 96-well plate following manufacturers’ instructions. Samples were run on a BD FACSCelesta (BD Biosciences) using the manufacturer-recommended settings for Legendplex bead acquisition. Data were analysed using the Legendplex v8 analysis software (Biolegend).

Gating was performed according to manufacturer guidelines.

### Untargeted metabolomics

Untargeted metabolomics of the caecal content was conducted on samples obtained from Figure 4.3A and carried out by MS-Omics as follows. Semi-polar metabolite profiling was performed by MS-Omics (Vedbæk, Denmark). The analysis was carried out using a Vanquish LC (Thermo Fisher Scientific) coupled to a Orbitrap Exploris 240 MS (Thermo Fisher Scientific). The UHPLC used an adapted method described by Doneanu et al. (UPLC/MS Monitoring of Water-Soluble Vitamin Bs in Cell Culture Media in Minutes, Water Application note 2011, 720004042en). An electrospray ionization interface was used as an ionization source. Analysis was performed in positive and negative ionization mode under polarity switching.

Untargeted data processing was performed using Compound Discoverer 3.3 (Thermo Fisher Scientific) and Skyline (25.1, MacCoss Lab Software) for peak picking and feature grouping, followed by an in-house annotation and curation pipeline written in MatLab (2022b, MathWorks). Identification of compounds were performed at four levels; Level 1: identification by retention times (compared against in-house authentic standards), accurate mass (with an accepted deviation of 3ppm), and MS/MS spectra, Level 2a: identification by retention times (compared against in-house authentic standards), accurate mass (with an accepted deviation of 3ppm). Level 2b: identification by accurate mass (with an accepted deviation of 3ppm), and MS/MS spectra, Level 3: identification by accurate mass alone (with an accepted deviation of 3ppm). Annotations on level 2b are based on accurate mass and MS/MS spectra measured with high resolution Orbitrap ESI-MS in mzCloud (Thermo Fisher Scientific), MassBank of North America (UC Davis) and the European MassBank (Helmholtz Centre for Environmental Research Leipzig). The annotations on level 3 are based on searches in the following libraries: Human metabolome database (version 5.0). Only metabolites identified at level 1 were used for the analysis conducted in this research.

### Targeted short chain fatty acid analysis

Short-chain fatty acid (SCFA) analysis was carried out by MS-Omics (Vedbæk, Denmark) using gas chromatography-mass spectrometry. Samples were acidified using hydrochloric acid, and deuterium-labeled internal standards were added. Analysis was performed in randomized order using a high-polarity column (Zebron™ ZB-FFAP, GC Cap. Column 15 m x 0.25 mm x 0.25 µm) installed in a GC (7890B, Agilent) coupled with a quadropole MS detector (5977B, Agilent). The system was controlled by ChemStation (Agilent). Peak areas were integrated using Skyline (25.1, MacCoss Lab Software), before quantification and curation using an in-house pipeline written in MATLAB (2022b, MathWorks). Matrix effects, carryover, noise levels, and precision were evaluated using corresponding quality control samples. In case of analytical overlap of SCFA peaks with matrix contaminants, peak deconvolution was performed using Skyline r skyline_version (MacCoss Lab Software).

### Metabolomics analysis

Data analysis on metabolomics was performed on centered log-ratio (CLR) transformed values to reduce compositionality bias and data skewness and perform standard statistical approaches (Aitchison et al., 2000). Principal component analysis (PCA) was performed on CLR-transformed values. Zeroes were replaced using the ‘const’ approach described by Lubbe and colleagues (Lubbe et al., 2021). Peak area values that fell below the limit of detection (LOD) were marked and replaced with the respective LOD value of that metabolite. The PERMANOVA implementation from the vegan library was used with 1000 permutations to find structural differences between control and treatment groups on a compositional level (Oksanen et al., 2012). To find features that were differentially abundant between groups, we fitted linear models for each CLR-transformed value. To correct for multiple tastings involved with the study and control for False Discovery Rates (FDR), features were selected based on the Benjamini-Hochberg adjustment procedure with a q-value threshold of 0.1. Metabolites and their log2fold change against controls were calculated and illustrated as volcano plots with FDR threshold. Of the differentially expressed metabolites that passed FDR correction, the top 40 were selected, and a literature search was conducted to identify the gut-microbial associated metabolites (total of 15; see **Table 1**). Metabolites that were significantly different between groups (FDR, q < 0.1) were used for enrichment analysis and inputted into the Metaboanalyst online pipeline (Chong et al., 2019) using the KEGG library as a reference. For Spearman’s correlation analysis, log-transformed peak area values of SCFAs were correlated against scaled breakpoint values per individual (**Table 2**).

### Quantification and statistical analysis

Data and graphs were analysed using RStudio (http://www.rstudio.com/) and Graphpad Prism (8.0.2). Technical outliers were detected using the Grubb’s test and removed prior to statistical analysis. Statistical tests are indicated in figure legends, and exact p values for comparisons are reported in Table S1 (main figures) and Table S2 (supplementary figures).

Unpaired t-tests were used to compare data between two unmatched groups. One-way ANOVA with Tukey’s multiple comparisons test was used to compare data between more than two unmatched groups. Two-way ANOVA was used to compare data between 2 categorically independent variables (e.g., SDV and ABX). Mixed effects models were used to assess differences between two or more groups across repeated measures, with Sidak’s multiple comparisons test. To assess within-subject comparisons mixed effects models with repeated measures and Dunnett’s multiple comparisons test was used. All tests were carried out at a significance level of p<0.05.

## Supplementary Information

**Figure S1.**
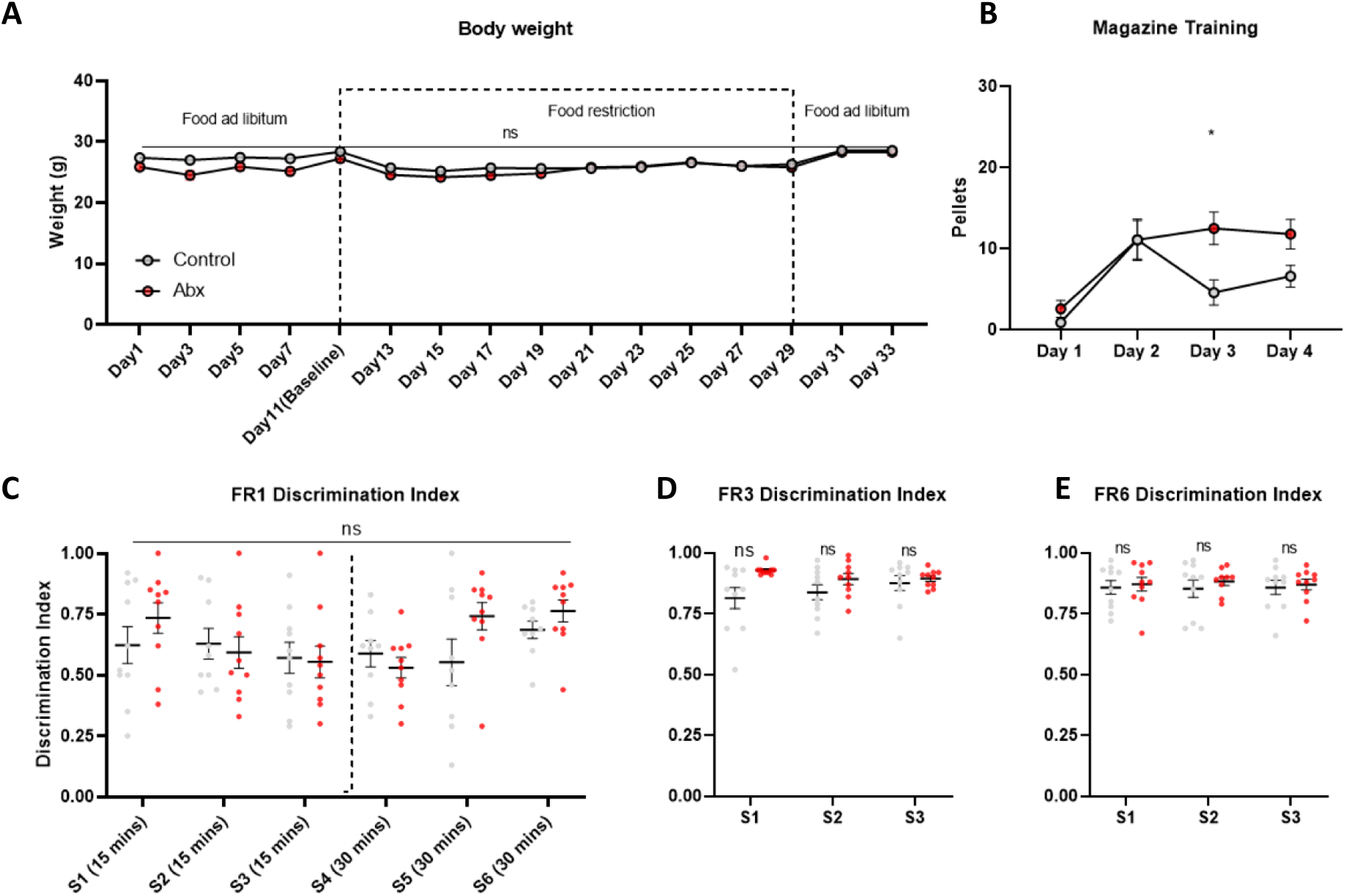
No significant effect of ABX on body weight or discrimination index throughout operant training protocol. Related to Figure 1. (A) No significant difference in mean body weights between ABX (n = 10) and control (n = 10) mice. (B) Freely available palatable reward consumption during magazine training. (C) No significant differences in mean discrimination index between ABX and control mice during the FR1 training protocol, (D) FR3 training protocol and (E) FR6 training protocol. Data presented as mean +/- SEM. Data analysed by mixed effects model with repeated measures followed by Sidak’s multiple comparisons test. * p<0.05, ** p<0.01, *** p <0.001 and **** p < 0.0001, ns = not significant. Detailed statistical information can be found in Table S2.

**Figure S2.**
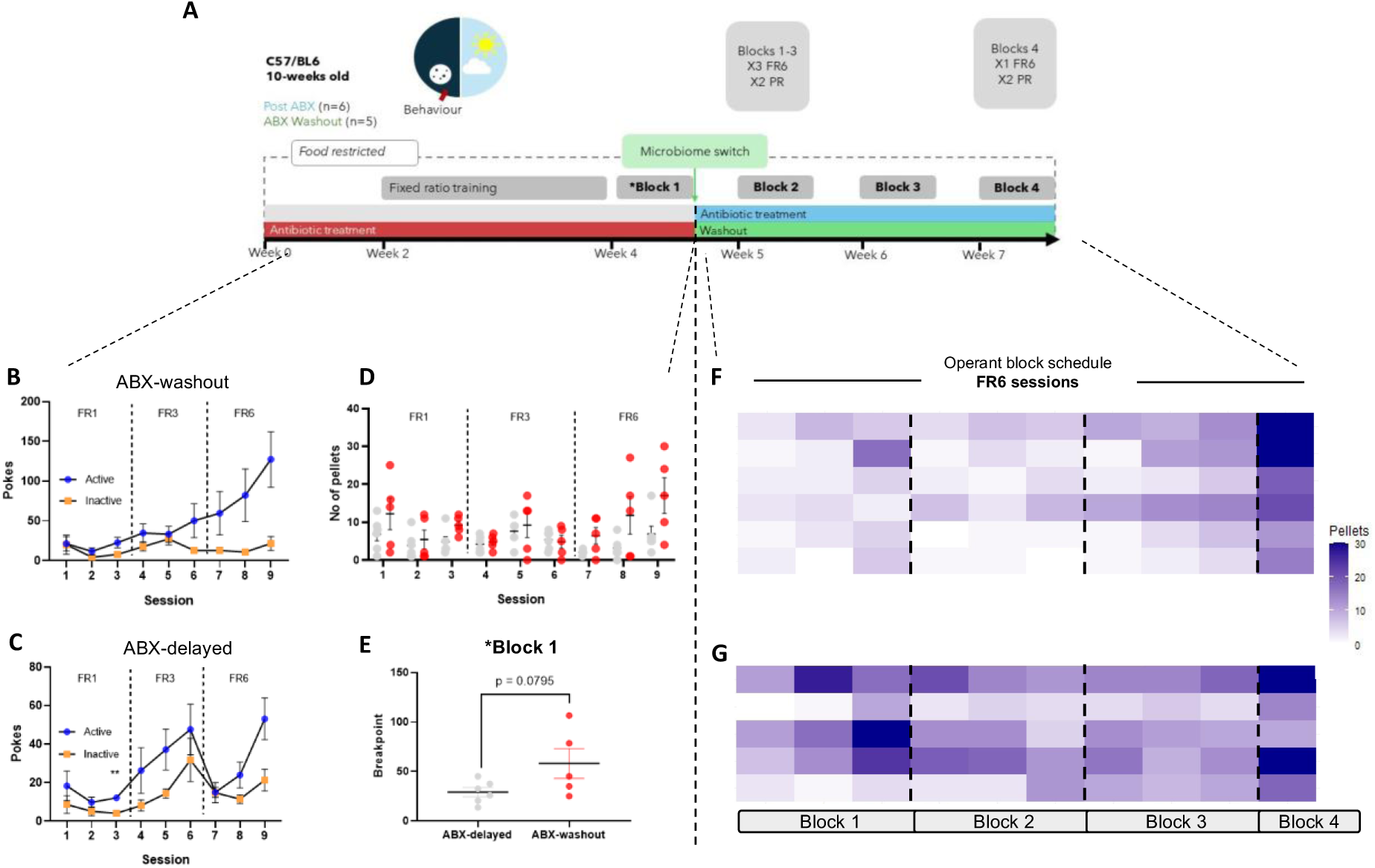
FR learning paradigm and FR6 sessions during block schedules of ABX-delayed and ABX-washout mice. Related to Figure 3. (A) Experimental timeline (B) Number of active and inactive nose pokes (mean +/- SEM) of ABX-washout mice (n = 5) across the FR1, FR3 and FR6 operant training sessions. (C) Active and inactive nose pokes (mean +/- SEM) of ABX-delayed mice (n = 6) across the FR1, FR3 and FR6 behavioural sessions. (F) Heatmap of within-subject number of pellets earned in FR6 sessions across 4 block operant schedules for pre- and post-ABX treatment (ABX-delayed group; n=6). (C) Heatmap of within-subject number of pellets earned in FR6 sessions across 4 block operant schedules for pre- and post-ABX-washout mice (ABX-washout group; n=5). Data presented as mean +/- standard error of mean (SEM). Mixed effects models with repeated measures followed by Sidak’s multiple comparisons test. PR breakpoints assessed via unpaired t test. * p<0.05, ** p<0.01, *** p <0.001, **** p < 0.0001 and ns = not significant. Detailed statistical information can be found in Table S2.

**Figure S3.**
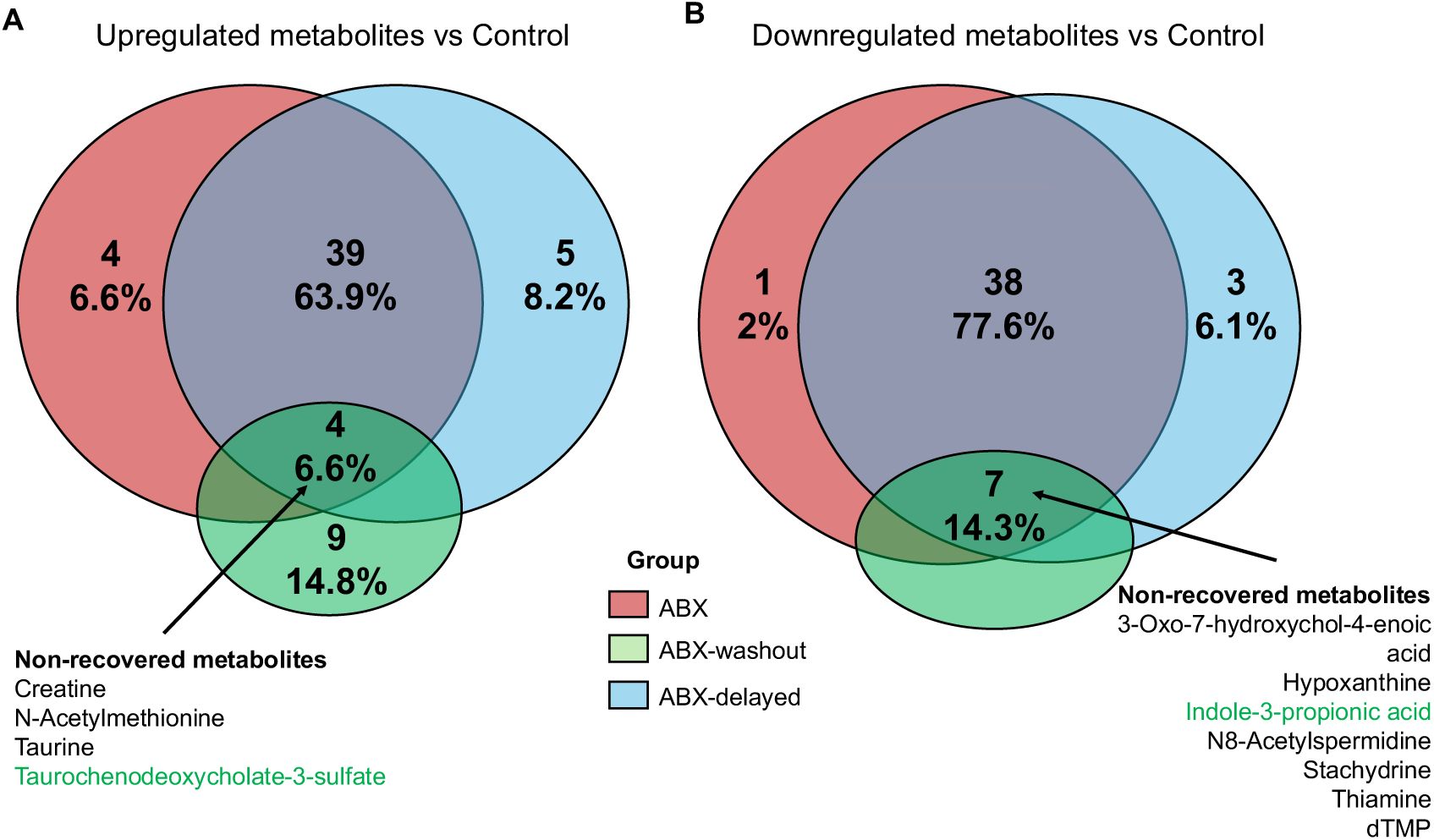
Overlapping metabolites differentially expressed against control across all groups from untargeted metabolomics. Related to Figure 4. (A, B) Venn diagrams illustrating (A) upregulated and (B) downregulated metabolites of ABX, ABX-washout and ABX-delayed against control groups (n= 6/ group). Arrow highlights metabolites that failed to recover.

**Figure S4.**
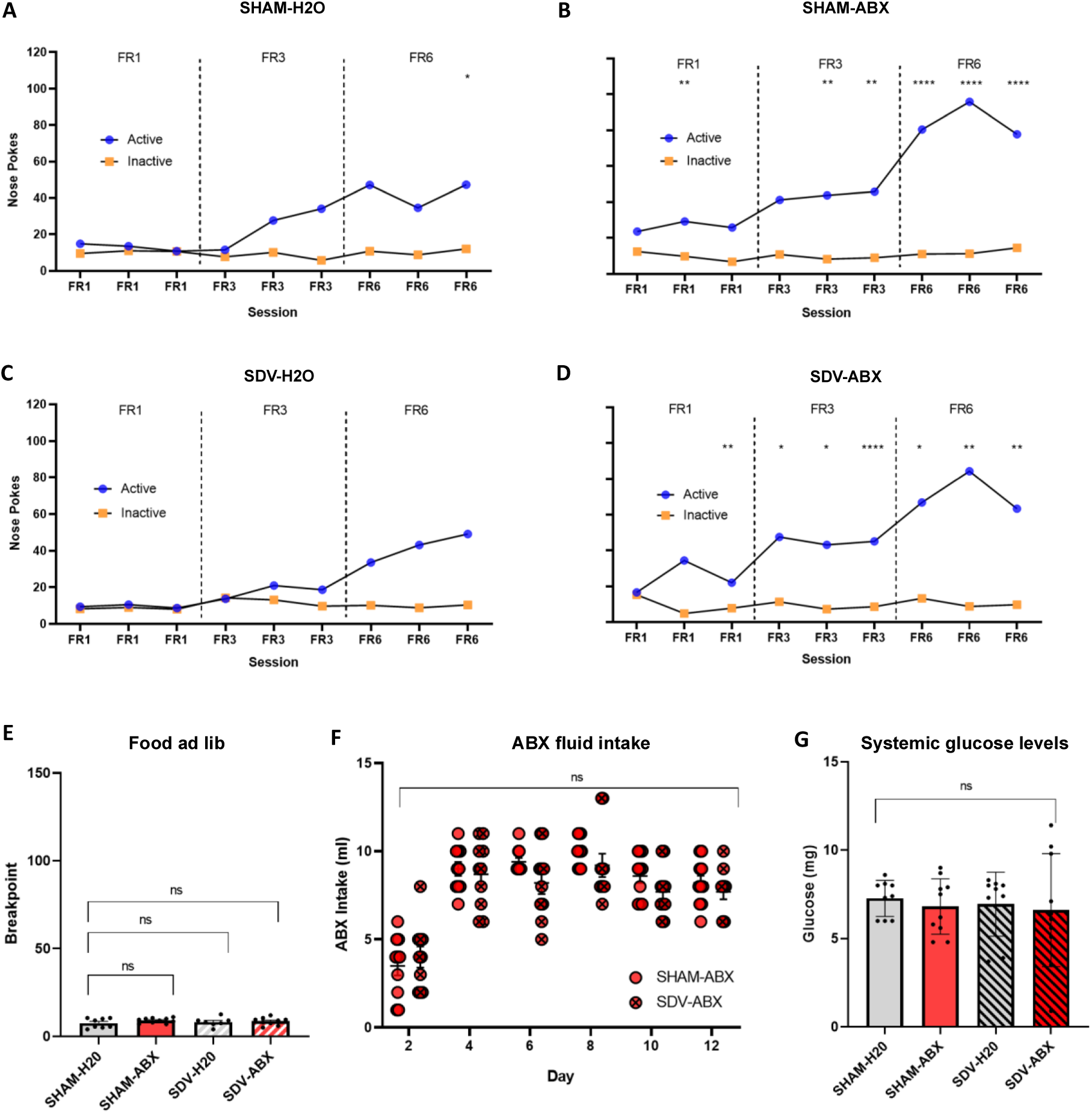
Additional operant behavioural data ABX fluid intake and systemic glucose readouts. Related to Figure 5. (A) Number of active and inactive nose pokes (mean +/- SEM) of SHAM-H2O mice across the FR1, FR3 and FR6 operant training sessions. (B) Active and inactive nose pokes (mean +/- SEM) of SHAM-ABX mice across the FR1, FR3 and FR6 behavioural sessions. (C) Number of active and inactive nose pokes (mean +/- SEM) of SDV-H2O mice across the FR1, FR3 and FR6 operant training sessions. (D) Active and inactive nose pokes (mean +/- SEM) of SDV-ABX mice across the FR1, FR3 and FR6 behavioural sessions. (A-D) Data analysed by mixed effects models with repeated measures followed by Sidak’s multiple comparisons test. (E) No significant effects of SDV or ABX on mean breakpoint when food is returned to ad libitum. Data analysed by two-way ANOVA (treatment x vagus) followed by Sidak’s multiple comparisons test. (F) No significant differences in mean ABX fluid consumption between SHAM-ABX and SDV-ABX. Data analysed by mixed effects models with repeated measures followed by Sidak’s multiple comparisons test. (G) No significant differences in systemic glucose levels between groups. Data analysed by two-way ANOVA (treatment x vagus) followed by Sidak’s multiple comparisons test. Data presented as mean +/- SEM. (E-G) SHAM-ABX n=10, SHAM-H2O n=8/9, SDV-H2O n=7/9, SDV-ABX n=9/10 successfully trained mice. * p<0.05, ** p<0.01, *** p <0.001, **** p < 0.0001 and ns = not significant. Detailed statistical information can be found in Table S2.

